# A Model of the Transition to Behavioral and Cognitive Modernity using Reflexively Autocatalytic Networks

**DOI:** 10.1101/2020.07.10.185512

**Authors:** Liane Gabora, Mike Steel

**Affiliations:** Department of Psychology, University of British Columbia, Okanagan Campus, Kelowna BC, Canada; Biomathematics Research Centre, University of Canterbury, Christchurch, New Zealand

**Keywords:** autocatalytic network, behavioral modernity, cognitive modernity, cultural evolution, Hohlenstein-Stadel Löwenmensch figurine, semantic network, Upper Paleolithic

## Abstract

This paper proposes a model of the cognitive mechanisms underlying the transition to behavioral and cognitive modernity in the Upper Paleolithic using autocatalytic networks. These networks have been used to model life’s origins. More recently, they have been applied to the emergence of *cognitive* structure capable of undergoing *cultural* evolution. Mental representations of knowledge and experiences play the role of catalytic molecules, the interactions among them (e.g., the forging of new associations or affordances) play the role of reactions, and thought processes are modeled as chains of these interactions. We posit that one or more genetic mutations may have allowed thought to be spontaneously tailored to the situation by modulating the degree of (1) divergence (versus convergence), (2) abstractness (versus concreteness), and (3) context-specificity. This culminated in persistent, unified autocatalytic semantic networks that bridged previously compartmentalized knowledge and experience. We explain the model using one of the oldest-known uncontested examples of figurative art: the carving of the Hohlenstein-Stadel Löwenmensch, or lion-man. The approach keeps track of where in a cultural lineage each innovation appears, and models cumulative change step by step. It paves the way for a broad scientific framework for the origins of both biological and cultural evolutionary processes.

## 1. Introduction

How did we become distinctively human? What enabled us to develop imagination, ingenuity, and complex belief systems? These questions are central to understanding who we are, how we got here, and where we are headed. Behavioral and cognitive modernity are thought to have come about between 100,000 and 30,000 years ago, as evidenced by the proliferation in cultural artifacts of both utilitarian and aesthetic value. (Although some researchers argue that the onset of behavioral modernity was less pronounced than once thought [70, 66, 94], and the concept of behavioral modernity itself has been called into question [23], this paper does not delve into these discussions so as to focus squarely on the task of modeling the cognitive changes underlying this cultural transition.) Some attribute this transition to an enhanced ability to process social information [99, 103]. Cognitive explanations have been proposed; for example, it has been attributed to the onset of conceptual fluidity [78], dual modes of information processing [29, 83], or enhanced working memory [20]. We propose that each of these proposals holds merit and that they are not mutually exclusive, but reflect the onset a new kind of semantic network structure, which is modeled here.

Although evidence of human culture dates back millions of years, behavioral-cognitive modernity is associated with the transition to cultural change that is not just adaptive (new innovations that yield some benefit for their bearers tend to predominate), but also cumulative (later innovations build on earlier ones), and open-ended (the space of possible innovations is not finite, since each innovation can give rise to spin-offs). In other words, culture became an *evolutionary* process [10, 15, 18, 26, 35, 52, 75, 88]. By *culture*, we mean extrasomatic adaptations, including behavior and artifacts, that are socially rather than genetically transmitted. Although cultural *transmission*—in which one individual acquires elements of culture from another—is observed in many species, cultural *evolution* is much rarer, and perhaps unique to our species.^1^

Critical to cultural evolution is the capacity to combine ideas, adapt existing solutions to new situations, and reframe information in one’s own terms.^2^ This paper uses network theory to address how this capacity arose. Networks allow for a comprehensive understanding of the dynamics of complex entities and their relationships [47]. Network-based approaches to characterizing the kind of cognitive structure that could sustain cultural evolution enable us to address the question of how minds carry out the contextual, combinatorial, and hierarchically structured thought processes needed to generate cumulative, adaptive, and open-ended cultural novelty [31, 40]. Here, to capture the self-organizing, self-maintaining, and indeed self-replicating nature of human cognition, rather than a generic semantic or neural network, we use an autocatalytic network. Autocatalytic network theory grew out of studies of the statistical properties of *random graphs* consisting of nodes randomly connected by edges [27]. As the ratio of edges to nodes increases, the size of the largest cluster increases, and the probability of a phase transition resulting in a single giant connected cluster also increases. The recognition that connected graphs exhibit phase transitions led to their application to efforts to develop a formal model of the origin of life (OOL), namely, of how abiogenic catalytic molecules crossed the threshold to the kind of collectively self-sustaining, self-replicating structure we call ‘alive’ [64, 63]. In the application of graph theory to the OOL, the nodes represent catalytic molecules and the edges represent reactions. It is exceedingly improbable that any catalytic molecule present in the primordial soup of Earth’s early atmosphere catalyzed its own formation. However, reactions generate new molecules that catalyze new reactions, and as the variety of molecules increases, the variety of reactions increases faster. As the ratio of reactions to molecules increases, the probability increases that the system will undergo a phase transition. When, for each molecule in a set, there is a catalytic pathway to its formation, the set is said to be collectively *autocatalytic*, and the process by which this state is achieved has been referred to as *autocatalytic closure* [64]. The molecules thereby become a self-sustaining, self-replicating structure (i.e., a living protocell [56]). Thus, the theory of autocatalytic networks has provided a promising avenue for modeling the OOL and thereby understanding how biological evolution began [105]. The approach is consistent with claims for the centrality of transitions across the life sciences [91].

Autocatalytic networks have been developed mathematically and generalized for cross-disciplinary application in other settings in the theory of Reflexively Autocatalytic Food set generated (RAF) networks [57, 93]. The term *reflexively* is used in its mathematical sense, meaning that every element is related to the whole. The term *food set* refers to the reactants that are initially present, as opposed to those that are the products of catalytic reactions. RAFs have been used extensively to model the origins of biological evolution [57, 93, 100, 105]. Thus, one strength of the approach is that by adapting a formalism that has been used successfully to model one evolutionary process to model another, we pave the way for a broad conceptual framework that can shed light on both [31, 5]. This is in keeping with the suggestion that autocatalytic networks may hold the key to understanding the origins of *any* evolutionary process, including the origin of culture [31, 32, 35, 38, 42].^3^ In application to culture, the products and reactants are not catalytic molecules but culturally transmittable *mental representations* ^4^ (MRs) of experiences, ideas, and chunks of knowledge, as well as more complex mental structures such as schemas and scripts (Tables 1 and 2).

**Table 1.** Application of graph theoretic concepts to the origin of life (OOL) and origin of culture (OOC).

| Graph Theory | Origin of Life (OOL) | Origin of Culture (OOC) |
| --- | --- | --- |
| node | catalytic molecule | mental representation (MR) |
| edge | reaction pathway | association |
| cluster | molecules connected via reactions | MRs connected via associations |
| connected graph | autocatalytic closure [63, 64] | conceptual closure<sup>5</sup> [31] |

**Table 2.** Abbreviations used throughout this paper.

| Abbreviation | Meaning |
| --- | --- |
| OOL | Origin of Life |
| OOC | Origin of Culture |
| MR | Mental Representation |
| RR | Representational Redescription |
| RAF | Reflexively Autocatalytic and Food set generated (F-generated) |
| CCP | Cognitive Catalytic Process |

Another strength of the approach is that because it distinguishes reactants that are external in origin—in our case, MRs that were acquired through social learning or individual learning of *existing* information—from those that are the products of internal reactions—in our case, MRs that come about through the creative generation of *new* information—MRs are tagged with their source. This enables us to model how networks emerge and to trace cumulative change in cultural lineages step by step.

In previous work, we used the RAF framework to model what is arguably the earliest significant transition in the archaeological record: the transition from Oldowan to Acheulean tool technology approximately 1.76 million years ago (mya) [42, 43]. We posited that this was precipitated by onset of the capacity for *representational redescription* (RR), in which the contents of working memory are recursively restructured by drawing upon similar or related ideas, or through concept combination. This enabled the forging of associations between MRs, and the emergence of hierarchically structured concepts, making it possible to shift between levels of abstraction as needed to carry out tasks composed of multiple subgoals. This culminated in what is referred to as a transient RAF, a critical step toward what has been referred to as *conceptual closure* [32], characterized by the emergence of persistent ‘autocatalytic’ cognitive structure. The application of RAFs presented in this paper builds on that work to model a pivotal cultural transition in human history that has been referred to as the “origins of art, religion, and science” [76]. We propose that behavioral and cognitive modernity was brought about by the emergence of an autocatalytic semantic network. We first summarize the archaeological evidence for a transition to behavioral modernity in the Upper Paleolithic. We then present our RAF model of the underlying cognitive transition that brought it about. Finally, we compare and contrast our proposal with existing literature.

## 2. Archaeological evidence for behavioral and cognitive modernity

We begin with a brief summary of the evidence for a transition to behavioral and cognitive modernity in the Upper Paleolithic.^6^ Although one can argue that the earliest stone tools marked the onset of a ‘proto’ form of cultural evolution, following the initial appearance of the Acheulian hand axe, the archaeological record exhibits considerable stasis^7^, and—with the exception of a more sophisticated knapping (Levallois) technique 200,000-400,000 years ago— little in the way of creative embellishment or improvement [97].

This changed in the Aurignacian period of the Upper Paleolithic, at which point there is evidence of recognizably human ways of living and thinking. The earliest evidence of behavioral and cognitive modernity comes from Africa less than 100,000 years ago, in Sub-Himalayan Asia and Australasia more than 50,000 years ago [80], and in Continental Europe until approximately 30,000 years ago [74]. This evidence consists of a proliferation of different complex, task-specific tools including effective cutting blades [3, 70]. It also marks the appearance of representational art [6, 82, 86], artifacts indicating personal symbolic ornamentation [25], complex living spaces [85], sophisticated ways of obtaining food, including aquatic resources [28], burial sites indicating ritual [58] and possibly religion [89]. The Upper Paleolithic is also widely believed to have marked the onset of modern syntactically rich language [12] (though some argue that language arose more gradually; c.f. [68]). In short, this period witnessed an unprecedented dramatic increase in the variety, utility, and aesthetic value of cultural artifacts.

A celebrated example of Upper Paleolithic art to which we will devote considerable attention is the Löwenmensch or ‘lion-man’ figurine from the Hohlenstein-Stadel cave in Germany (Figure 1). This figurine, carbon-dated to the Interpleniglacial period between 35,000 and 40,000 years ago, is one of the oldest-known zoomorphic (animal-shaped) sculpture in the world, and one of the oldest-known examples of figurative art. It measures 31.1 cm, and was carved out of mammoth ivory using a flint stone knife.

**Figure 1.**
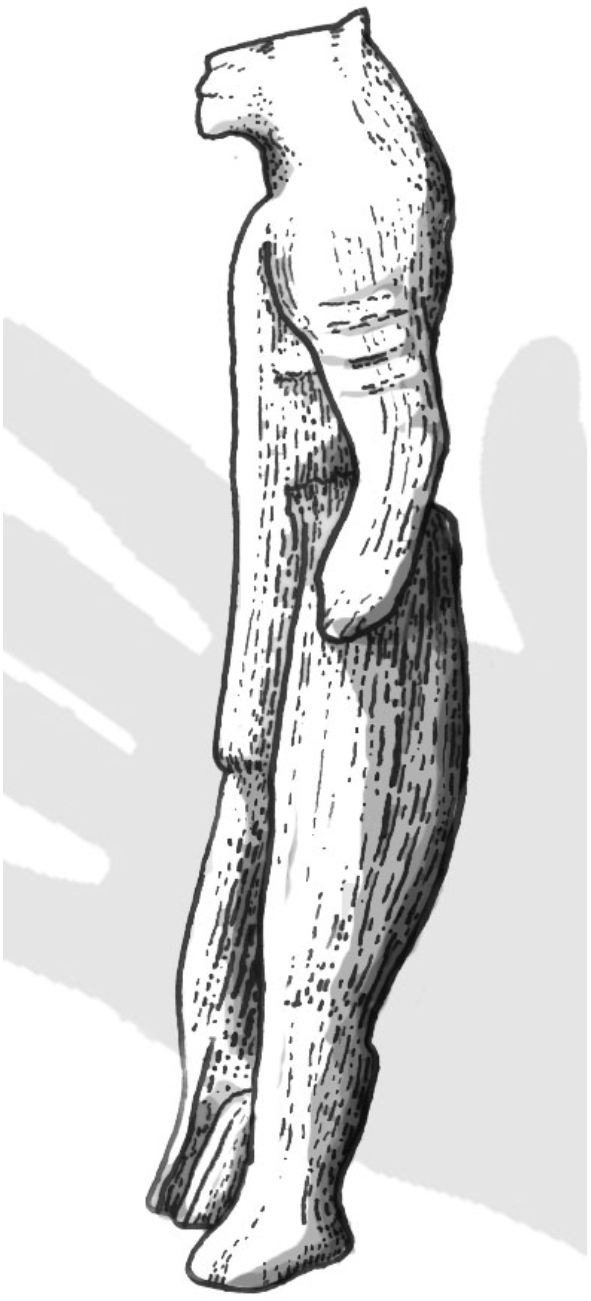
Sketch of the Löwenmensch or ‘lion-man’ fgurine from the Hohlenstein-Stadel cave in Germany. According to the Ulm Museum, ^14^C dates put it at an age of 35,000 to 40,000 years. The hand indicates its relative size (Obtained with permission from the artist, Cameron Smith).

## 3. An underlying cognitive transition

The model of the transition to cognitive and behavioral modernity in the Upper Paleolithic developed here grew out of the hypothesis that it was due to the onset of *contextual focus:* the capacity to, in a spontaneous and ongoing manner, shift between convergent and divergent modes of thought, thereby tailoring ones’ mode of thought to one’s situation [33, 34, 40]. Focused attention is conducive to *convergent thought* because the activation of neural cell assemblies is constrained enough to zero in on the most defining properties. In this compact form, the contents of thought are more readily amenable to deliberate executive level operations. In convergent thought, one can access only close associates of the current thought: items that are highly related to it with respect to the most conventional, default context. For example, FIG and PLUM are close associates because they are both fruits, and they are most commonly thought about with respect to their membership in the category FRUIT.

In contrast, defocused attention is conducive to divergent thought because it causes diffuse activation of neural cell assemblies in memory, such that obscure (but potentially relevant) properties come into play [32, 34, 36, 37]. This is useful for creative tasks, and when one is in need of a new approach or innovative solution. Divergent thought may include more details of the current subject of thought, or incorporate related items; one is simply taking in more of the situation and its associations. In divergent thought, one can access remote associates: words or concepts that are not related to each other with respect to their most conventional, default context. Highly divergent thought may result in cross-domain thinking, in which ideas from different domains are combined, or a solution to a problem in one domain is borrowed from another domain.

Although divergent thought is useful for escaping local minima, it confers the risk of getting perpetually side-tracked, whereby irrelevant thoughts readily intrude, interfering with survival tasks. Unless the capacity for divergent thought is accompanied by the ability to reign it in, it would be counterproductive. Therefore, it seems likely that in the pre-modern mind, before the advent of the capacity to shift along the spectrum from convergent to divergent, all mental contents were processed convergently, such that each successive thought was a close associate of its predecessor, and remote associates were not accessed.

Contextual focus came about through the onset of the capacity to adjust the focus of attention to current constraints and affordances, making it more focused or diffuse, as needed, thereby stretching or shrinking conceptual space, and tailoring working memory to task demands (or lack thereof, as in mind wandering). Contextual focus made it possible to shift between (1) convergent thought to modify the content of working memory on the basis of close associates when that was sufficient, and (2) divergent thought to usher in remote associates when ‘stuck in a rut’. The theory that contextual focus can have a transformative impact on cultural evolution was tested using an agent-based model [40]. Incorporating the ability to shift between convergent and divergent processing modes into neural network-based agents in the agent-based model resulted in an increase in the mean fitness of cultural outputs.

The model that follows builds on the hypothesis that behavioral modernity was due to the onset of contextual focus, but goes further in positing that thought acquired the capacity to shift along a multimodal spectrum through spontaneous tuning of the following three variables.

### 3.1. Divergence

The first variable is the capacity to shift between divergent and convergent thought, as discussed above.

### 3.2. Level of abstraction

The second variable is *degree of abstraction*. It has been shown that there is what is called a *basic level* of abstraction (e.g., BIRD, as opposed to ANIMAL or SPARROW) that mirrors the correlational structure of properties in the object’s real-world perception and use [90]. Categories form, and are first learned and perceived, at this basic level, before they are further discriminated at the subordinate level (e.g., SPARROW), and abstracted at the superordinate level (e.g., ANIMAL).^8^ Since basic-level categories contain the degree of abstraction most useful for carrying out daily activities [90], it seems reasonable that they precede other levels of abstraction not just developmentally but evolutionarily. We posit that the arrival of behavioral/cognitive modernity involved onset of the capacity to shift along the hierarchy from abstract to concrete, thereby identifying relatedness at different hierarchical levels, and incorporating these distinctions into one’s mental model of the world. Abstraction provides another means of connecting MRs, but instead of forging a remote association between them, it makes explicit that they are both instances of some more general MR (e.g., LION and MAMMOTH are both instances of ANIMAL). Thus, the second variable involves the capacity to shift from basic-level categories to other levels of abstraction.

### 3.3. Context-specificity

To generate ideas and solutions that are not just new but also task-relevant may require thought that is not just divergent but also context-specific [36]. Thus, the third variable is *context-specificity*: the degree to which thought is biased by a specific motivating contextual factor such as a goal or desire [8]. Divergent thought need not *always* be context-specific (e.g., during mind-wandering or writing free verse). However, context-specific divergent thought allows one to access information that is related to the current contents of working memory in ways that may be unconventional yet precisely relevant to the current situation [73]. For example, thinking of lions in the context of wishing for an inspirational reminder of a lion’s power might prompt one to modify one’s concept of lion to incorporate the possibility of carving a lion. This unusual context makes this remote yet feasible relationship ‘pop out’.

### 3.4 Multimodality

The spectrum of thought is multimodal, where by a ‘mode’ we refer to a particular combination of these three variables (e.g., divergent, abstract, and context-specific). In short, we posit that by using this multimodal spectrum to modify how one thought gives way to the next, cognitive processes could be carried out more effectively. Moreover, the fruits of one mode of thought could become ingredients for another mode, thereby facilitating the forging of a richly integrated network of understandings about the world and one’s place in it, sometimes referred to as a *worldview* [32, 38]. This, we posit, set the stage for behavioral modernity.^9^

Note that although we may be ‘wired for culture’ [69], and the cognitive changes underlying this cultural transition may have (directly or indirectly; see [4]) involved one or more genetic mutations [33, 41, 20, 21], we are not proposing that control over these variables came online instantaneously, nor that control over each of them arose simultaneously. The challenge may have been not so much to *possess* the capacity to change these variables as to *coordinate* them so as to continuously tune one’s mode of thought in response to changing task demands and effectively navigate semantic space. The evolutionary and developmental tinkering required to achieve this could explain the lag between anatomically modern *Homo sapiens* 200,000 to 100,000 years ago, and behavioral modernity 100,000 to 30,000 ago.

## 4. Autocatalytic networks (RAFs)

The mathematical model we will describe and analyse in this paper is based the notion of RAFs. Our use of RAFs as an underlying model is based on three considerations: (i) the model has a high degree of generality, which has allowed its application to explain transition events in a variety of fields (as mentioned above), (ii) it has a well-developed mathematical theory, and (iii) in earlier work [42, 43] RAFs have provided a way to investigate cognitive processes (and transitions in them in early cultural evolution) which we develop further here in a more complete mathematical model.^10^ Thus, in order to explain our approach we first summarize the key concepts of RAF theory.

A *catalytic reaction system* (CRS) is a tuple 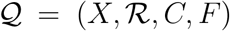 consisting of a set *X* of molecule types, a set 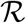 of reactions, a catalysis set *C* indicating which molecule types catalyze which reactions, and a subset *F* of *X* called the food set. A *Reflexively Autocatalytic and F-generated* set—i.e., a RAF—is a non-empty subset 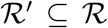 of reactions that satisfies the following two properties:

1. *Reflexively autocatalytic*: each reaction 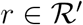 is catalyzed by at least one molecule type that is either produced by 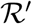 or is present in the food set *F*; and
2. *F-generated*: all reactants in 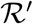 can be generated from the food set *F* via a series of reactions only from 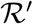 itself.

A set of reactions that forms a RAF is simultaneously self-sustaining (by the *F*-generated condition) and (collectively) autocatalytic (by the RA condition; as each of its reactions is atalyzed by a molecule associated with the RAF). A CRS need not have a RAF, but when it does there is a unique maximal one. Moreover, a CRS, may contain many RAFs, and it is this feature that allows RAFs to evolve, as demonstrated (both in theory and in simulation studies) through selective proliferation and drift acting on possible subRAFs of the maxRAF [57, 100]. In the OOL context, a RAF emerges in systems of polymers (molecules consisting of repeated units called monomers) when the complexity of these polymers (as measured by their maximum length) reaches a certain threshold [64, 79]. The phase transition from no RAF to a RAF incorporating most or all of the molecules depends on (1) the probability of any one polymer catalyzing a given reaction that forms another polymer, and (2) the maximum length (number of monomers) of polymers in the system. This transition has been formalized and analyzed (mathematically and via simulations), and applied to real biochemical systems [53, 54, 55, 57, 79], ecology [19], and cognition [42, 43]. RAF theory has proven useful for identifying how phase transitions might occur, and at what parameter values.

### 4.1 Terminology

We now introduce the mathematical framework and terminology that will be used to model the transition to cognitive modernity. All mental representations (MRs) in a given individual *i* are denoted *X*_*i*_, and a particular MR *x* = *x*_*i*_ in *X*_*i*_ is denoted by writing *x* ∈ *X*_*i*_. As in an OOL RAF, we have a *food set*; for individual *i*, this is denoted *F*_*i*_. In the origin of culture (OOC) context, *F*_*i*_ encompasses MRs for individual *i* that are either innate, or that result from direct experience in the world, including natural, artificial, and social stimuli. *F*_*i*_ includes everything in the long-term memory of individual *i* that was not the direct result of individual *i* engaging in RR. This includes information obtained through social learning from *someone else* who may have obtained it by way of RR. For example, if individual *i* learns from individual *j* how to edge a blank flake through percussive action, this is an instance of social learning, and the concept EDGING is therefore a member of *F*_*i*_.

*F*_*i*_ also includes existing information obtained by *i* through individual learning (which, as stated earlier, involves learning from the environment by nonsocial means), so long as this information retains the form in which it was originally perceived (and does not undergo redescription or restructuring through abstract thought). The crucial distinction between food set and non-food set items is not whether another person was involved, nor whether the MR was originally obtained through abstract thought (by *someone*), but whether the abstract thought process originated in the mind of the individual *i* in question. Thus, *F*_*i*_ has two components:

- *S*_*i*_ denotes the set of MRs arising through direct stimulus experience that have been encoded in individual *i*’s memory. It includes MRs obtained through social learning from the communication of an MR *x*_*j*_ by another individual *j*, denoted S_*i*_[*x*_*j*_], and MRs obtained through individual learning, denoted S_*i*_[*l*], as well as contents of memory arising through direct perception that does not involve learning, denoted S_*i*_[*p*].
- *I*_*i*_ denotes any *innate knowledge* with which individual *i* is born.

A particular catalytic event (i.e., a single instance of RR) in a stream of abstract thought in individual *i* is referred to as a *reaction,* and denoted 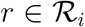. A stream of abstract thought, involving the generation of representations that go beyond what has been directly observed, is modeled as a sequence of catalytic events. Following [42], we refer to this as a *cognitive catalytic process* (CCP). The set of reactions that can be catalyzed by a given MR *x* in individual *i* is denoted *C*_*i*_[*x*]. The entire set of MRs either *undergoing* or *resulting from r* is denoted *A* or *B*, respectively, and a member of the set of MRs undergoing or resulting from reaction *r* is denoted *a* ∈ *A* or *b* ∈ *B*.

The term *food set derived*, denoted ¬*F*_*i*_, refers to mental contents that are *not* part of *F*_*i*_ (i.e. ¬*F*_*i*_ consists of all the products *b* ∈ *B* of all reactions *r* ∈ *R*_*i*_). In particular, ¬*F*_*i*_ includes the products of any reactions derived from *F*_*i*_ and encoded in individual *i*’s memory. Its contents come about through mental operations *by the individual in question* on the food set; in other words, food set derived items are the direct product of RR. Thus, ¬*F*_*i*_ includes everything in long-term memory that *was* the result of one’s own CCPs. ¬*F*_*i*_ may include a MR in which social learning played a role, so long as the most recent modification to this MR was a catalytic event (i.e., it involved RR).^11^

The set of *all* possible reactions in individual *i* is denoted 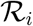. The mental contents of the mind, including all MRs and all RR events, is denoted 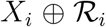. This includes *F*_*i*_ and ¬*F*_*i*_. Recall that the set of all MRs in individual *i*, including both the food set and elements derived from that food set, is denoted *X*_*i*_.

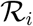 and *C*_*i*_ are not prescribed in advance; because *C*_*i*_ includes remindings and associations on the basis of one or more shared property, different CCPs can occur through interactions amongst MRs. Nevertheless, it makes perfect mathematical sense to talk about 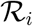 and *C*_*i*_ as sets. Table 3 summarizes the terminology and correspondences between the OOL and the OOC.

**Table 3.** Terminology and correspondences between the Origin of Life (OOL) and the Origin of Culture (OOC).

| Term | Origin of Life (OOL) | Origin of Culture (OOC) |
| --- | --- | --- |
| $X_i$ | all molecule types in protocell $i$ | all mental representations (MRs) in individual $i$ |
| $x \in X_i$ | a molecule in $X_i$ | a MR in $X_i$ |
| $F_i$ | food set for protocell $i$ | innate or directly experienced MRs by $i$ |
| $r \in \mathcal{R}_i$ | a particular reaction in $i$ | a particular representational redescription (RR) in $i$ |
| $C_i[x]$ | reactions catalyzed by $x$ in $i$ | RR events ‘catalyzed’ by $x$ in $i$ |
| $(x, r) \in C$ | $x$ catalyzes $r$ | $x$ ‘catalyzes’ redescription by $r$ |
| $a \in A$ | member of set of reactants in $r$ | member of set of MRs undergoing $r$ |
| $b \in B$ | member of set of products of $r$ | member of set of MRs resulting from $r$ |
| $\neg F_i$ | non food set for $i$ (i.e., all $B$ of $\mathcal{R}_i$) | MRs resulting from $R_i$ (i.e., all $B$ of $R_i$) |

Our model includes elements of cognition that have no obvious parallel in the OOL. We denote the subject of attention at time *t* as 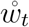. It may be an external stimulus, or a MR retrieved from memory. Any other contents of 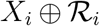 that are accessible to working memory, such as close associates of 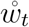, or recently attended MRs, are denoted *W*_*t*_, with *W*_*t*_ constituting a very small subset of 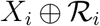. The focus here is on how non-food set derived MRs (i.e., a non-empty ¬*F*) emerge and connect giving rise to a semantic network that is reflexively autocatalytic and food set generated.

## 5. RAF model of the cognitive transition

We now use the RAF formalism to model the transition to behavioral/cognitive modernity in the Upper Paleolithic. To address how the mind as a whole acquired autocatalytic structure, the model is, by necessity, abstract. It does not distinguish between semantic memory (memory of words, concepts, propositions, and world knowledge) and episodic memory (personal experiences); indeed, we are sympathetic to the view that these are not as distinct as once thought [67]. Nor does it address how MRs are obtained (i.e., whether through Hebbian learning versus probabilistic inference). Although MRs are represented simply as points in an *N*-dimensional space (where *N* is the number of distinguishable differences, i.e., ways in which MRs could differ), our model is consistent with models that use convolution [60], random indexing [61], or other methods of representing MRs.

We assume that associations form between MRs but do not address whether they are due to similarity or co-occurrence, or whether they are learned through Bayesian inference [45] or other means. We view associations as probabilistic; when we say that an association was forged between two MRs we mean a spike in the probability of one MR evoking another, which we refer to as the ‘catalysis’ of one MR by the other. We view context as anything external (e.g., an object or person) or internal (e.g., other MRs) that influences the instantiation of a MR in working memory. Although our approach is influenced by how context is modeled in quantum approaches to concepts [1, 2], it is not committed to any formal approach to modeling context. MRs are composed of one or more *concepts:* mental constructs such as CAT or FREEDOM that enable us to interpret new situations in terms of similar previous ones. The rationale for treating MRs as catalysts comes from the literature on concept combination, which provides extensive evidence that when concepts act as contexts for each other, their meanings change in ways that are often non-trivial and defy classical logic [1, 2, 49, 84]. The extent to which the meaning of one MR is modified by another is referred to here as its *reactivity*. A given MR’s reactivity varies depending on the other MRs present in working memory.^12^. Although we do not explicitly model the dimensionality of semantic space itself (i.e., the features or properties of MRs), we do so indirectly, by representing hierarchical structure in terms of reactivity, as explained below. Our model hinges on the fact that interactions between two or more MRs in working memory alter (however slightly) the network of association strengths [16, 71]. Conceptual closure is achieved and a cognitive RAF network emerges when, for each MR, there is an associative pathway to its formation; in other words, any given concept can be explained using other concepts, and new ideas can be re-framed in terms of existing ones.

We now show how the RAF framework is used to model the emergence of a persistent and integrated cognitive RAF, through onset of the capacity to spontaneously control the ‘spectrum of thought’ variables introduced in Section Three, and summarized in Table 4. In this table, the variables *γ*_*D*_, *γ*_*A*_, and *γ*_*C*_ quantify the three variables: divergence (*D*), abstractness (*A*) and context-specificity (*C*), respectively.

**Table 4.** Examples of the three variables of the spectrum of thought.

| Variable | Example | Symbol |
| --- | --- | --- |
| divergence | LION → KILL → POWER | $\gamma_D$ |
| abstractness | LION → ANIMAL → ANIMATE BEING | $\gamma_A$ |
| context-specificity | LION (context: desire to possess lion’s power) → LION FIGURINE | $\gamma_C$ |

We model the capacity to shift between convergent and divergent thought by introducing a metric geometry. We let *d* denote the *semantic distance* between an item *m* in memory *M*_*t*_ and an item in working memory 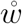. In convergent thought, the semantic distance *d* between successive contents of working memory remains small, and only close associates catalyze RR reactions and participate in CCPs. In contrast, by spontaneously engaging in divergent thought when stymied, or as a form of mental exploration or mind-wandering, the modern mind gained access to remote associates (i.e., items for which the semantic distance *d* to the content of working memory was large). These remote associates catalyzed RR reactions, and participated in CCPs. Thus, divergent thought could (in our terminology) bring about reactions amongst previously unconnected MRs, including MRs from different knowledge domains. The variable *γ*_*D*_ determines how ‘remote’ an associate can be in order to catalyze an update (i.e., how ‘far afield’ one looks for ingredients for one’s stream of thought). Thus, *γ*_*D*_ provides a threshold on *d* that increases as one shifts from convergent to divergent thought.

The more abstract a concept, the more associations it can have with other MRs. Therefore, we represent hierarchical semantic structure from concrete instances to increasingly abstract concepts in terms of reactivity. Consequently, the more abstract (as opposed to concrete) *x* in individual *i* is, the larger the value of *C*_*i*_[*x*]. Thus, abstract concepts facilitate the navigation of semantic space through CCPs. For example, during the transition from thinking about a particular sharp axe to thinking about the abstract quality of sharpness, abstractness increases, and therefore so does the reactivity, potentially leading the CCP to something quite different from a sharp axe, such as a ‘claw’.

As mentioned earlier, the capacity for divergent thought could be made even more useful by broadening the sphere of associates in a context-specific manner, such that one’s current needs bias the retrieval of information from memory. This facilitates the forging of new connections between MRs that would be irrelevant in most contexts but are relevant in the current one. We make the notion of context-specificity more precise by introducing a context-dependent association structure. As above, we let *d* denote the *semantic distance* between an item *m* in memory *M*_*t*_ and an item in working memory 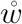. A small value of 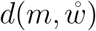 means that in the current context, *m* is closely related to 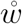 whereas a large value of 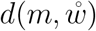 means that with respect to the current context, they are distantly related.

Let C_*t*_ denote a context at time *t* (which is determined by the goals and needs of the individual at time *t*). We can represent the context-dependent associations explicitly by writing 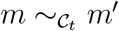 if *m* and *m*’ are related with respect to context C_*t*_.

For *m* to catalyze a cognitive updating reaction

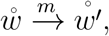

*m* should satisfy at least one of the following two properties, where *γ*_*D*_ is as described above, and *γ*_*C*_ is the extent to which context can facilitate the catalysis of a particular RR reaction:

i. 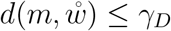, or
ii. 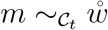 and 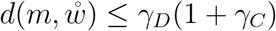.

In other words, for *m* to catalyse a RR reaction involving 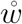, either the default semantic distance to *m* must be sufficiently small that divergent thought makes it accessible, or it is pulled within reach because context-specificity warps semantic space in such a way as to make this particular association salient. In addition, a particular context C_*t*_ at time *t* may mean that a stimulus *s*_*t*_ that is relevant to the current contents of working memory, catalyzes a RR reaction 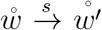 that would not occur otherwise. For example, seeing an animal puncture a food source with its claw could be a source of ideas for how to make a sharper tool.

### 5.1. Example: The Hohlenstein-Stadel figurine

We now make the transition from premodern to modern mind more concrete using the example of the Löwenmensch or ‘lion-man’ figurine from the Hohlenstein-Stadel cave, discussed in Section 2. Although we cannot know exactly how the Hohlenstein-Stadel figurine was created, by reverse-engineering the process it is possible to infer what conceptual structure would, at a minimum, have had to be in place [39, 98, 102]. We carry this out using available evidence, such as our knowledge that the lion was the largest and most dangerous predator in the ecosystem of the Interpleniglacial [87, 65], and likely a source of fear and awe due to its power and aggression [48]. Since the word ‘representation’ is often used to refer to an internal, mental construct of something in the world, to avoid confusion, we use the term *iconic* to refer to an object that represents something else in a way that is not merely symbolic but captures its physical attributes.

We now consider the sequence of steps culminating in the creation of the lion man, summarized in Table 5 and depicted in Figures 2 and 3. Note that the steps culminating in the Hohlenstein-Stadel figurine, were preceded by, and dependent upon, the development of lithic reduction (i.e., knapping and carving) techniques. (These are not discussed here, since they are the subject of another paper [43].) Note that the role of catalysts tends to be played by MRs that represent needs or questions, since like catalysts these speed up conceptual change that would otherwise occur very slowly, yet their participation in this process does not fundamentally change them (that is, the degree to which the need is satisfied may change, but the mental representation of the need does not).

**Table 5.** The sequence of steps culminating in the creation of the Hohlenstein-Stadel Löwenmensch figurine.

| Step | Description | Origin | Mode |
| --- | --- | --- | --- |
| 1. | Carve from stone → carve (from something) | RR | abstract |
| 2. | Transmission of lithic reduction techniques | social learning | convergent |
| 3. | Carve functional tool → carve (something) | RR | divergent, abstract |
| 4. | Transmission of CARVE (something) | social learning | convergent |
| 5. | Carve (something) → carve iconic likeness | RR | divergent, concrete |
| 6. | Combine human form and lion head →<br>internalize lion's power | RR | divergent, cross-domain,<br>context-specific |
| 7. | Assimilate features of lion and man | individual learning | divergent |
| 8. | Carve Löwenmensch | individual learning + RR | divergent |

**Figure 2.**
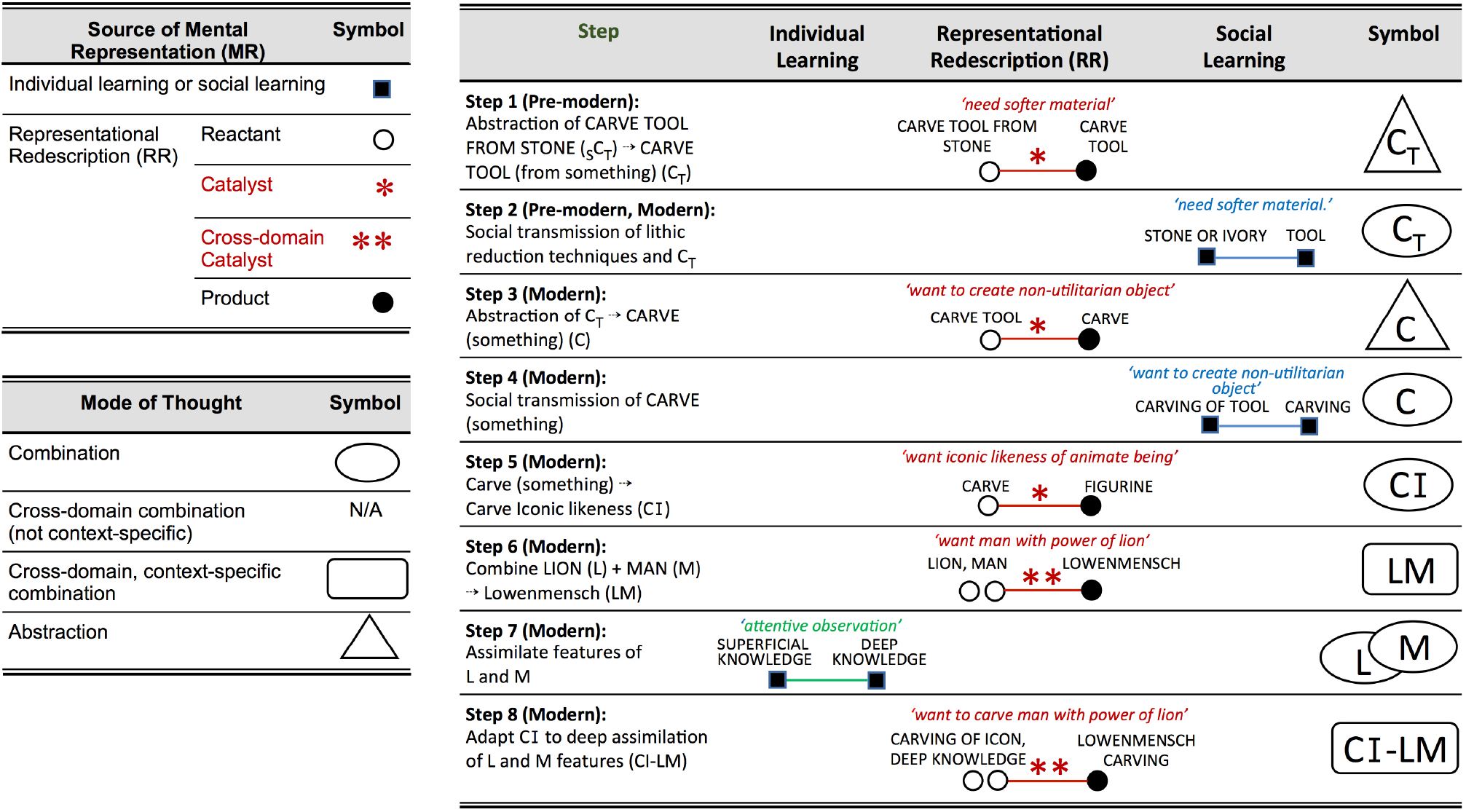
Top left: Sources of mental representations involved in the creation of the Hohlenstein-Stadel figurine, and the symbols used to depict them. Bottom left: Modes of thought and symbols used to depict them. Right: Steps involved in the creation of the figurine. Top two rows show the steps that occurred prior to the Upper Paleolithic; subsequent rows depict steps that took place during the Upper Paleolithic.

**Figure 3.**
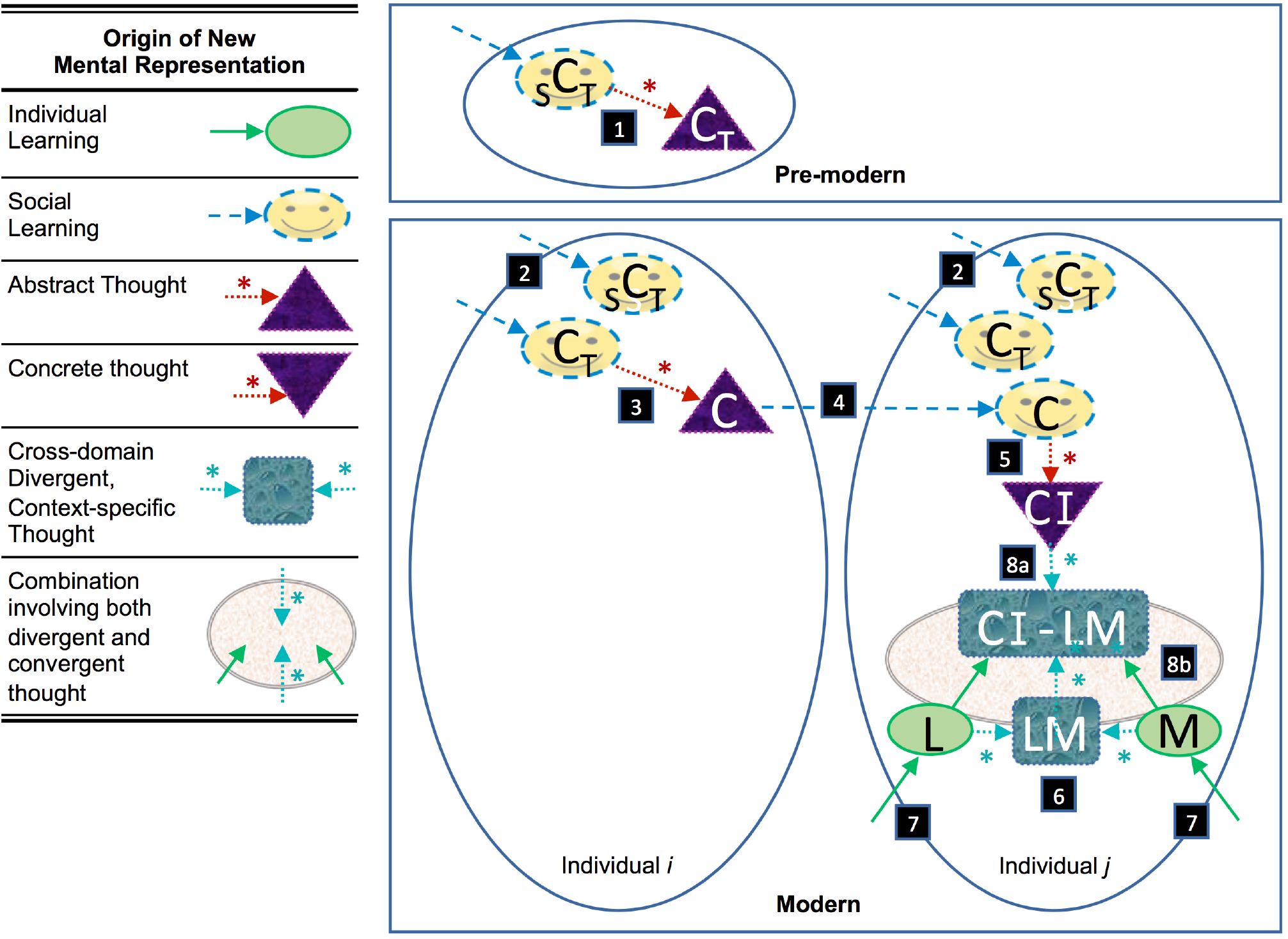
Steps culminating in creation of Hohlenstein-Stadel figurine. Meanings of symbols are defined in Figure 2 (see text for details).

1. **Form abstract concept, CARVE** (from any suitable material). This consisted of abstracting the general concept of lithic reduction with stone as the source material to lithic reduction using any suitable material (e.g., mammoth ivory). There is evidence that the capacity to abstract a general concept from particular instances dates back to at least 1.76 mya (well before the Upper Paleolithic) [95].
2. **Social learning of first step.** Creative contributions to culture begin with a preparation stage involving thorough assimilation of relevant background knowledge [106]. Thus, the first step that took place in the Upper Paleolithic involved the social learning of existing knapping and carving techniques, including the abstract concept CARVE (from any suitable material).
3. **Abstraction of CARVE TOOL to CARVE (something)**. The next step was to extricate the concept CARVE TOOL from its conventional function of generating something utilitarian such as a hand axe. This resulted in a the abstract concept CARVE, which could now be applied in domains other than technology, such as art. This would have likely involved divergent, abstract thought. The existence of objects in bone, ochre, and ostrich eggshell with geometric engravings from southern Africa, dates this to at least 77,000 years ago [50], with earlier dates for ivory engraving in China [44].
4. **Social learning of the abstract concept CARVE (something)**. The carver of the Hohlenstein-Stadel figurine acquired the abstract concept CARVE (something) through social learning.^13^
5. **Apply CARVE (something) to the domain of figures (animal and human), yielding CARVE FIGURINE.** We may never know exactly what motivated the first artisan who took the step of carving an iconic likeness, a figurine. It may have been the product of idle mind wandering. An alternative and perhaps more likely explanation is that it was shaped by a goal or desire, such as to (1) know the depicted subject more deeply, or (2) gain a sense of control or mastery over it, or (3) preserve a memory of it, or (4) have constant access to a feeling associated with it, such as the feeling of power associated with a lion. Whatever the motive, it involves taking the concept CARVE and applying it to a new domain, that of ANIMATE BEINGS.
6. **Combine LION HEAD with HUMAN BODY.** We also do not know what motivated this step. Like the previous step, it is possible that it was the product of idle mind wandering. It could be that by endowing a human body like ours with the head of a lion, the artisan hoped that those who held it would internalize the lion’s power as their own. An alternative possibility is that it held some religious significance. Again, for the purpose of this model it is not essential to know which of these is correct, for whatever the underlying motive, this cross-domain combination would have required RR using divergent, context-specific thought.
7. **Assimilate features.** To carve an iconic figurine, knowledge of lithic reduction techniques is not sufficient; the artisan would have had to deeply absorb the physical characteristics of lions and humans through individual learning. We characterize this process as divergent because it involves assimilating the details and, potentially, any feelings they evoke. The artisan would then have used RR to creatively adapt this technical knowledge to the new task of rendering a figure in ivory.
8. **Carve figurine.** The actual carving of the figurine would have required RR in divergent mode to creatively adapt known carving techniques to the new task of rendering the detailed characteristics of the lion and human forms. Engagement in a tactile process meant that thought was concrete, ensuring that the features of the figurine were recognizably human or lion-like.

The entire mental trajectory through the spectrum of thought culminating in the creation of the Hohlenstein-Stadel figurine is depicted in Figure 4.

**Figure 4.**
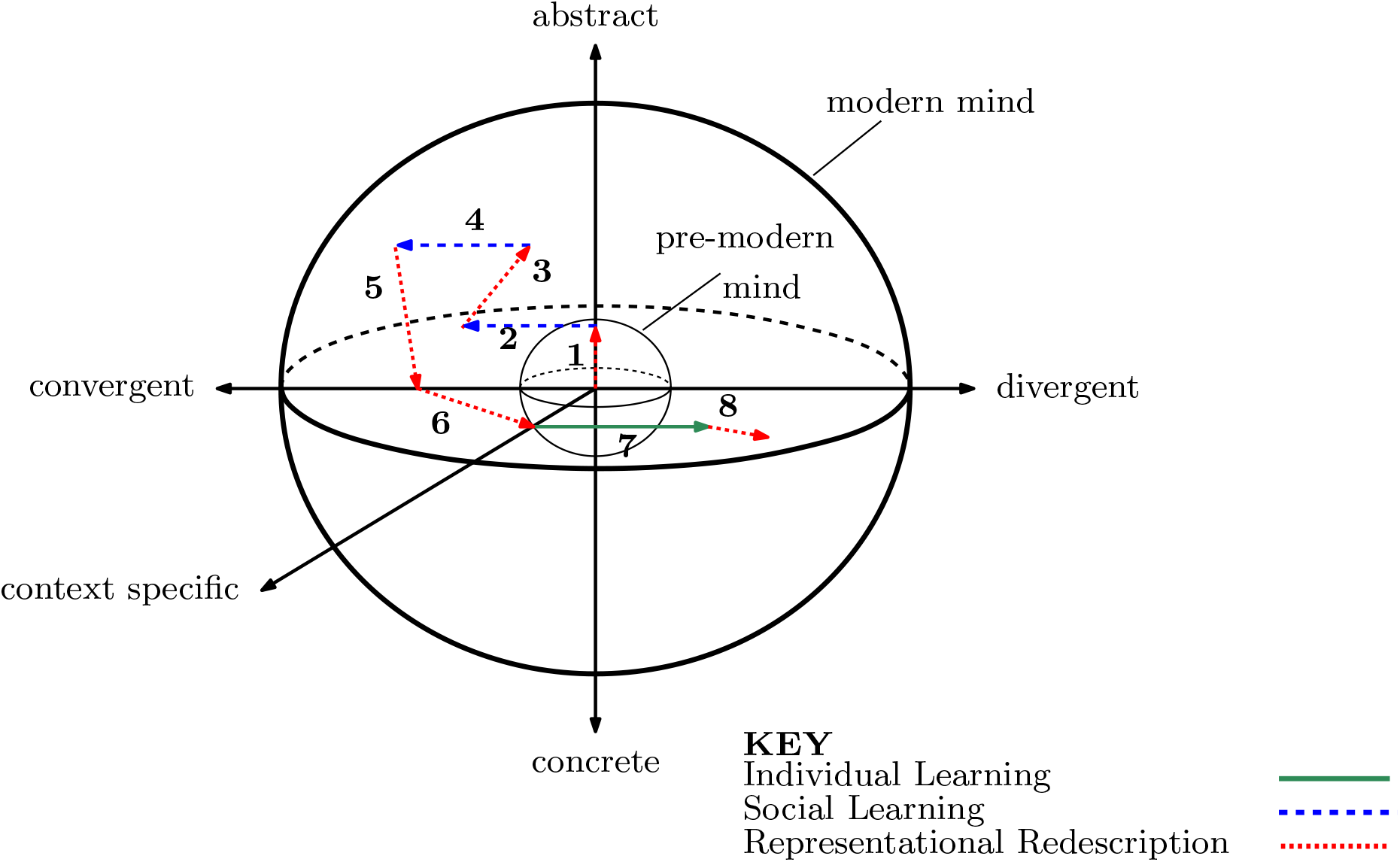
Trajectory through the spectrum of thought culminating in the Hohlenstein-Stadel figurine. Convergent-to-divergent is on the *x*-axis, abstractto-concrete is on the *y*-axis, and degree of context-specificity is on the *z*-axis. Different combinations of these three variables comprise different ‘modes’ of thought (i.e., ways of navigating memory and processing information). Although the premodern mind could, to some degree, form abstractions, its thought trajectories used only a tiny portion of this space, as indicated by the small sphere. It was therefore restricted to a single mode of thought. The modern mind could engage in all combinations of these three variables, thereby engaging in many modes of thought, as indicated by the large sphere. The numbered arrows correspond to the eight steps listed in Table 5; thus, they depict how the mode of thought shifted over course of the figurine-making process.

## 6. The transition to cognitive modernity

In the pre-modern mind, information is thought to have been compartmentalized into domain-specific modules, and RR only operated within particular domains of human inquiry (e.g., ‘tools’) [78]. Pre-modern cognition was largely (though not entirely) restricted to basic level categories—an intermediate level of abstraction—without the context-specific RR needed for cross-domain thinking. This was modeled by restricting RAFs to closed subsets of MRs that only reacted with other members of the same subset, resulting in RAF structure that was transient and fragmented [42, 43].

We now describe a simple mathematical model of the formation and persistence of cognitive RAFs culminating in behavioral/cognitive modernity and the cultural transition of the Upper Paleolithic. Let 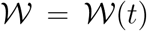 be a continuous measure of the scope (or content) of working memory of an individual at time *t* (where the continuous variable *t* varies over the lifetime of that individual). Cognitive processes make use of items obtained through individual learning, social learning, or RR, with items persisting in working memory for a short (but variable) time. As in [42], we model this as a non-deterministic process. Let 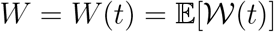 denote the expected (i.e., mean) value of 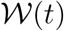. A direct implementation of the dynamical model in [42] then leads to the following nonlinear first-order equation:

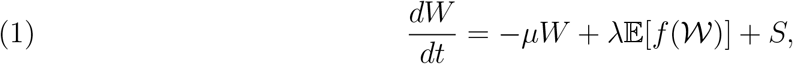

where *S* = *S*(*t*) ≥ 0 is a measure of information that is externally derived, either through individual learning or social learning at time *t*, the value 1*/μ* is the mean time that items remain in working memory, *λ* describes the rate of RR reactions, and *f* is a certain (unspecified) function that satisfies only the minimal requirements that *f* (0) = 0, *f*’(0) *>* 0, and *f* is concave (this last assumption recognises that 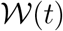 is bounded). Simple default choices for *f* would be *f* (*x*) = min{*x, K*} or *f* (*x*) = *x*(1 − *x/K*), though we do not explicitly assume either of these here.

Crucially, the parameter *λ* also depends on total memory (a richer memory of knowledge and experiences allows more opportunity to catalyse RR reactions) and it is influenced by the three variables described above (*γ*_*D*_, *γ*_*A*_, *γ*_*C*_) that we propose distinguish pre-modern from modern cognition. More precisely, if 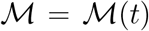 denotes a continuous measure of the scope of total memory at time *t*, and 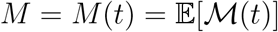 is the expected (mean) value of 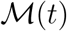, then *λ* is dependent on *M* (i.e. *λ* = *λ*(*M*)). Thus, Equation (1) is coupled to the growth in *M*, and a simple model for the dynamics of expected total memory *M* is the first-order linear differential equation:

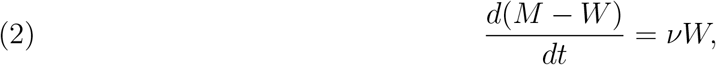

where *ν* ∈ (0, 1) parameterises the extent to which items in working memory become encoded in long-term memory (which may also depend on time, as *ν* may vary during the lifetime of the individual). The coupled (non-linear) system of Equations (1) and (2) leads to certain predictions. In particular, when *λ* lies below a critical threshold (dependent on the other parameters), CCPs do not form or persist, and thoughts are driven externally (individual learning or social learning). However, once *λ* passes this threshold, CCPs can form and persist indefinitely, even when the term *S* (in Eqn. (1)) drops to zero. The justification of these two claims and further mathematical details are provided in the Appendix.

The ability to shift between convergent and divergent thought, to consider the same item at multiple levels of abstraction, and to allow context to bias retrieval from memory by adjusting *γ*_*D*_, *γ*_*A*_ and *γ*_*C*_, provide distinct and complementary mechanisms for *λ* to change. If the resulting new MRs are encoded in long-term memory, the positive dependence of *λ* on *M* provides more routes for catalysis of CCPs. This increases *M* − *W* from Eqn. (2), which, in turn, influences the dynamics of *W* by Eqn. (1).

The modern mind could carry out logical operations during convergent thought, and make new connections using divergent thought. Divergent thought could be biased toward a specific need by making thought more contextual. The modern mind could also shift up and down the hierarchy from concrete to abstract. By tuning the mode of thought along the three variables of the above multimodal spectrum to match the situation one is in, the modern mind acquired the capacity to work out how elements of their world were interrelated, and where each element fitted with respect to the whole (i.e., the integrated internal model of the world, or *worldview*). The modern mind could now synthesize different domains of understanding into a coherent web of understandings, using not only basic level concepts [90] but also higher or lower levels of abstraction, from fine-grained details to the ‘big picture’, as appropriate.

The worldview of the modern mind is a ‘metabolism’ in the sense that it has in place entropy-defying processes that maintain its organization. Like the protocell that constituted the earliest structure that could be said to be alive, the structure of the autocatalytic cognitive network as a whole is now maintained through the interactions amongst its parts. New experiences are interpreted, understood, and encoded in memory, in terms of existing cognitive structure already in place; thus, the ‘small-world’ structure of the memory expands.

## 7. Comparison with other theories

This model builds on the theory that the burst of creativity in the Paleolithic was due to the onset of contextual focus: the capacity to shift between divergent and convergent modes of thought [33]. That theory is superficially similar to the proposal that the distinguishing feature of human cognition is our capacity for dual processing [29, 83].^14^ Our model builds on both the contextual focus and dual processing theories by positing that a single-variable spectrum of thought is insufficient to achieve an integrated internal model of the world.

Our model is consistent with Mithen’s [77] theory that the transition was due to the connecting of domain-specific information processing modules, thereby enabling metaphorical thinking and cognitive fluidity: the capacity to combine ideas from different domains, fuse different knowledge processing techniques, or adapt a solution to one problem to a different problem. It is also consistent with Coolidge and Wynn’s [20] theory that it was due to expanded working memory.^15^ Conceptual fluidity and expanded working memory are underwritten by divergent thought but, as explained above, the capacity to engage in divergent thought without the capacity to control *how* divergent one’s thinking is, would be perilous. Although it is not the focus of this paper, like [20], as well as [21] we are sympathetic with the view that genetic mutation was involved (see [41]).

Our proposal is consistent with the view that complex languages, symbolic representation, and myth lay at the heart of this transition [13, 14, 24]. However, we put the emergence of a persistent (i.e., stable) and integrated RAF network as central, with language both facilitating and being facilitated by this structure. Given evidence of recursive reasoning well before behavioral modernity, our framework is inconsistent with the hypothesis that the onset of recursive thought enabled mental time travel and cognitive modernity [22, 96]; nevertheless, the ability to shift through a multimodal spectrum of thought would have brought on the capacity to make vastly better use of it. The proposal that behavioral modernity arose due to onset of the capacity to model the contents of other minds, sometimes referred to as the ‘Theory of Mind’ [99], is somewhat underwritten by recursive RR, since the mechanism that allows for recursion is required for modeling the contents of other minds (though in this case the emphasis is on the social impact of recursion, rather than the capacity for recursion itself). Our proposal is also consistent with explanations for behavioral modernity that emphasize social-ecological factors [30, 103], but places these explanations in a broader framework by suggesting a mechanism that aided not just social skills but other skills (e.g., technological) as well.

## 8. Discussion and conclusions

Formal models exist of many aspects of human cognition, such as learning, memory, planning, and concept combination. However, there is little in the way of formal models of how they came to function together as an integrated whole, and how the unique cognitive abilities of *Homo sapiens* came about. RAF networks provide a means of addressing these questions. Building on earlier models of the cognitive transition underlying the earliest origins of human culture and the invention of the Acheulean hand axe, resulting in a transient autocatalytic structure, in this paper, we developed a model of the transition to a persistent, integrated RAF network. We proposed that rapid cultural change in the Middle-Upper Paleolithic required the ability to, not just recursively redescribe the contents of thought, but also tailor the ‘reactivity’ of thought to the current situation. This was accomplished through continuous, spontaneous tuning of three variables that concern not the content of thought *per se,* but how it is processed. The first involves shifting between convergent and divergent processing. The second involves shifting between concrete and abstract representations. The third involves biasing divergent processing according to a pressing need or context. Together, these enabled *Homo sapiens* to reflect on the contents of thought from different perspectives and at different levels of abstraction. This culminated in the crossing of a threshold to conceptual closure and the achievement of self-organizing autocatalytic semantic networks that spanned different knowledge domains, and routinely integrated new information by reframing it in terms of current understandings.

The model is highly simplified, and we do not know the precise details of the cognitive events modelled here took place (though the model does not hinge on these details). We hope that future research will incorporate inhibition (in conjunction with the existing catalysis), as well as a more sophisticated representation of the interactions amongst MRs [1, 2] and a dynamic representation of context [59, 101]. A platform for the computational modeling of RAFs exists (https://github.com/husonlab/catlynet), and we hope to apply it to cognitive RAFs. There remains much work to be done on how cognitive RAFs replicate and evolve (see [5] for informal suggestions in this regard) and on the developmental question of how persistent, integrated RAF networks emerge in the mind of a child.

We also hope that future research will build on the direction taken here by comparing the cognitive RAF model with other standard semantic network models [9, 7, 11, 62, 72]. Although these standard semantic networks suffice for modeling semantic structure in individuals, we believe that the RAF approach will turn out to be superior because it distinguishes semantic structure arising through social or individual learning (modeled as food set items) from semantic structure *derived from* this pre-existing material (modeled as non-food set items generated through abstract thought processes that play the role of catalyzed reactions). This makes it feasible to model how cognitive structure emerges, and to trace lineages of cumulative cultural change step by step. It also frames this project within the overarching scientific enterprise of understanding how evolutionary processes (be they biological or cultural) begin, and unfold over time.

## Authors’ contribution

L.G. provided the conceptual framework and wrote the majority of the text; M.S. contributed mathematical modelling details, including the formal proofs in the Appendix.

## Acknowledgments

We thank the two anonymous reviewers for several helpful suggestions and additional references for an earlier version of this manuscript. We also thank Dietrich Radel for assistance with the formatting of references.

## Data accessibility

This article has no additional data.

## 9. Appendix Mathematical details and justification of predictions based on Equations (1) and (2).

In the following arguments, we treat *λ* as a constant over the short time-frame considered in the dynamics of CCPs, since the dependence of *λ* on *M* applies over considerably longer time-scales. Moreover, in treating *λ* as a constant and setting *S* = 0, Eqn (1) is technically not an ordinary differential equation (i.e. it is not of the form Φ(*W, dW/dt*) = *S*(*t*)), since 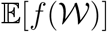 is not, in general, a function of *W*. For example, for *f* (*x*) = *x*(1 − *x/K*), Eqn (1) becomes:

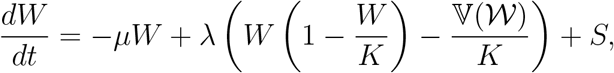

where 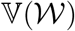 is the variance of 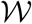 at time *t*. Note also that the dynamics of 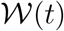 is not determined by the behaviour of *W* (*t*); the latter just represents the expected (average) value of the former.

Turning to the first prediction of this model, observe that:

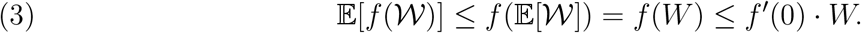

The first inequality in (3) is by Jensen’s inequality for the concave function *f* (see e.g. [46]). The second inequality also uses the concavity of *f* together with the conditions *f* (0) = 0 and *f*’(0) *>* 0. Thus if *λ < μ/f*’(0), we have:

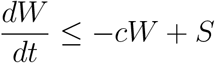

for *c* = (*μ* − *λf*’(0)) *>* 0. Consequently, once *S* declines to zero, so too does *W*, and by the Markov inequality (see e.g. [46]), we have (for any *E* > 0):

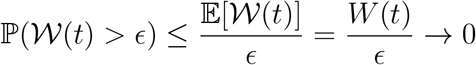

as *t* increases, which establishes the first prediction.

Now suppose that *λ > μ/f*’(0). Select *η >* 0 sufficient small so that *β* := (*μ* + *η*)*/λ < f*’(0). By the concavity of *f*, it follows that, for some *γ* ≥ 0, we have:

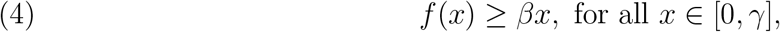

since the line *y* = *βx* and the function *y* = *f* (*x*) both pass through the origin; however, the latter function has a strictly greater slope at the origin.

Now suppose that 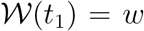, where *w* ∈ (0*, γ*). Considering the process moving forward from time *t*_1_, with the initial condition W(*t*_1_) = *w* we then have at *t* = *t*_1_:

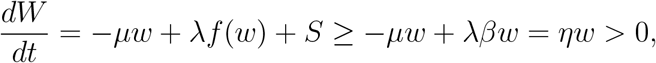

where the first inequality is from (4) together with *S*(*t*_1_) ≥ 0. In summary, when *λ* passes above the threshold *μ/f*’(0) and 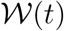 is small but non-zero the expected value of 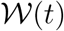 begins to increase, due to CCPs in working memory (and even with *S* = 0).

## Footnotes

1 The term ‘cultural evolution’ is occasionally used in a less restricted sense to refer to the generation and transmission of novelty without the requirement of cumulative, adaptive, open-ended change (e.g., [104]).

2 Although some attribute cultural evolution uniquely to an increase in the *number* of ideas and cognitive skills (or ‘cognitive gadgets’) [51], not their interactivity.

3 For related approaches, see [5, 17, 81].

4 Although we use the term ‘mental representation’, our model is consistent with the view (common amongst ecological psychologists and in the situated cognition and quantum cognition communities) that what we call mental representations do not ‘represent’, but instead act as contextually elicited bridges between mind and world.

6 A more detailed discussion can be found elsewhere [40, 41].

7 Note, however, that we cannot know the extent to which lack of evidence of cultural embellishment in the early record is due to taphonomic biases—i.e., biases in what gets preserved over time (such as lack of preservation of softer materials) [4].

8 Note that abstract processing is not the same as convergent processing. An item at a particular level of abstraction, such as LION, would, in convergent thought, be held in working memory in a compact manner stripped of details, whereas during divergent thought, it would be rich in the characteristics of, and feelings evoked by, lions. One might speculate that richly detailed visions of religious deities occurs in a mode of thought that is abstract yet divergent.

9 Note that, in this view, language enhanced not just the ability to communicate and collaborate (thereby accelerating the pace of cultural innovation), but also the ability to think ideas through for oneself and manipulate them in a manner that was controlled, deliberate, and multimodal.

10 The question of whether the underlying reality being modelled is precisely described by a RAF is more subtle, and beyond the scope of this paper.

11 1This distinction between food set and food set-derived may not be so black and white as portrayed here but for simplicity we avoid that subtlety for now.

12 For example, in a study of the influence of context and mode of thought on the perceived meanings of concepts (as measured by property applicabilities and exemplar typicalities), the concept PYLON was rated low as an exemplar of HAT; however, in the context FUNNY (as in ‘worn to be funny’), it was rated high as an exemplar of HAT [101]. Thus, the degree to which PYLON qualified as an instance of a HAT changed depending on the context. The context FUNNY had an even greater effect on the rating of MEDICINE HAT (as in the name of the Canadian town) as an instance of HAT. We say that the *reactivity* was high here because the context exerted a dramatic influence on the perceived meaning of the concept HAT

13 We cannot know for certain that it was not invented independently (particularly given the distance between Hohlenstein-Stadel and southern Africa).

14 Dual processing posits that humans engage in not just a primitive implicit Type 1 mode for free association and fast ‘gut responses’, but also an explicit Type 2 mode for deliberate analysis. However, although dual processing makes the split between older, more automatic processes and newer, more deliberate processes, contextual focus theory posits that pre-modern thought was intermediate between two extremes (each valuable in different ways): a divergent mode based on relationships of correlation, and a convergent mode based on relationships of causation. Earlier hominids’ memories were coarser-grained, so there were fewer routes for meaningful associations, and less processing of previous experiences. Rather than convergent or divergent processing of previously assimilated material, there was greater tendency to focus on the here and now, so items in memory tended to remain in the same form as when they were originally assimilated. For a comparison of the divergent thought and dual processing theories see [92].

15 Working memory is just the part of memory that is, at any moment, working.

## References

[1] D. Aerts, J. Broekaert, L. Gabora, and S. Sozzo, Generalizing prototype theory: A formal quantum framework, Frontiers in Psychology (Cognition), 7 (2016), p. 418, https://doi.org/10.3389/fpsyg.2016.00418.

[2] D. Aerts, L. Gabora, and S. Sozzo, Concepts and their dynamics: A quantum theoretical model, Topics in Cognitive Science, 5 (2013), pp. 737–772, https://doi.org/10.1111/tops.12042.

[3] S. Ambrose, Paleolithic technology and human evolution, Science, 291 (2001), pp. 1748–1753.

[4] C. Andersson, A. Törnberg, and P. Törnberg, An evolutionary developmental approach to cultural evolution, Current Anthropology, 55 (2014), pp. 154–174.

[5] C. Andersson and P. Törnberg, Toward a macroevolutionary theory of human evolution: The social protocell, Biological Theory, 14 (2019), pp. 86–102, https://doi.org/10.1007/s13752-018-0313-y.

[6] M. Aubert, P. Setiawan, A. Oktaviana, M. Brumm, P. H. Sulistyarto, E. W. Saptomo, and et al., Palaeolithic cave art in borneo, Nature, 564 (2018), pp. 254–257.

[7] A. Baronchelli, R. Ferrer-i-Cancho, R. Pastor-Satorras, N. Chater, and M. H. Christiansen, Networks in cognitive science., Trends in Cognitive Sciences, 17 (2013), pp. 348–360, https://doi.org/10.1016/j.tics.2013.04.010.

[8] L. W. Barsalou, Context-independent and context-dependent information in concepts, Memory & cognition, 10 (1982), pp. 82–93.

[9] R. E. Beaty, M. Benedek, P. J. Silvia, and D. L. Schacter, Creative cognition and brain network dynamics, Trends in Cognitive Science, 20 (2016), pp. 87–95.

[10] R. A. Bentley, M. W. Hahn, and S. J. Shennan, Random drift and culture change, Proceedings of the Royal Society of London. Series B: Biological Sciences, 271 (2004), pp. 1443–1450, https://doi.org/10.1098/rspb.2004.2746.

[11] R. F. Betzel and D. S. Bassett, Generative models for network neuroscience: Prospects and promise, Journal of The Royal Society Interface, 14 (2017), p. 20170623.

[12] D. Bickerton, Language evolution: A brief guide for linguists, Lingua, 117 (2007), pp. 510–526.

[13] D. Bickerton, More than nature needs: Language, mind and evolution, Harvard University Press, Cambridge, 2014.

[14] D. Bickerton and E. Szathmáry, Biological foundations and origin of syntax, MIT Press, Cambridge, 2009.

[15] R. Boyd and P. Richerson, Culture and the evolutionary process, University of Chicago Press, Chicago, 1988.

[16] J. Brockmeier, After the archive: Remapping memory, Culture and Psychology, 16 (2010), pp. 5–35, https://doi.org/10.1177/1354067X09353212.

[17] K. R. Cabell and J. Valsiner, The catalyzing mind: Beyond models of causality (Annals of Theoretical Psychology, Volume 11), Springer, Berlin, 2013, https://doi.org/10.1007/978-1-4614-8821-7.

[18] L. L. Cavalli-Sforza and M. W. Feldman, Cultural transmission and evolution: A quantitative approach, Princeton University Press, Princeton, NJ, 1981.

[19] R. Cazzolla Gatti, B. Fath, W. Hordijk, S. Kauffman, and R. Ulanowicz, Niche emergence as an autocatalytic process in the evolution of ecosystems, Journal of Theoretical Biology, 454 (2018), pp. 110–117, https://doi.org/10.1016/j.jtbi.2018.05.038.

[20] F. L. Coolidge and T. Wynn, Working memory, its executive functions, and the emergence of modern thinking, Cambridge Archaeological Journal, 15 (2005), pp. 5–26.

[21] M. Corballis, The origins of modernity: Was autonomous speech the critical factor?, Psychological Review, 111 (2004), pp. 543–552, https://doi.org/10.1037/0033-295X.111.2.543.

[22] M. Corballis, The recursive mind: The origins of human language, thought and civilization, Princeton University Press, Princeton, NJ, 2011, https://doi.org/10.2307/j.ctt6wpzjd.

[23] S. Davies, Behavioral modernity in retrospect, Topoi, (2019), pp. 1–12.

[24] T. Deacon, The symbolic species: The coevolution of language and the brain, Norton, New York, 1997.

[25] F. D’Errico, N. Bartond, A. Bouzouggar, H. Mienis, D. Richter, and P. Lozouet, Ad-ditional evidence on the use of personal ornaments in the Middle Paleolithic of North Africa, Proceedings of the National Academy of Sciences, USA, 106 (2009), pp. 16051–16056, https://doi.org/10.1073/pnas.0903532106.

[26] M. Enquist, S. Ghirlanda, and K. Eriksson, Modelling the evolution and diversity of cumulative culture, Philosophical Transactions of the Royal Society B: Biological Sciences, 366 (2011), pp. 412–423, https://doi.org/10.1098/rstb.2010.0132.

[27] P. Erdös and A. Rényi, On the evolution of random graphs, Publication of the Mathematical Institute of the Hungarian Academy of Sciences, 5 (1960), pp. 17–61.

[28] J. M. Erlandson, The archaeology of aquatic adaptations: Paradigms for a new millennium, Journal of Archaeological Research, 9 (2001), pp. 287–350.

[29] J. Evans, Dual-process accounts of reasoning, judgment and social cognition, Annual Review of Psychology, 59 (2008), pp. 255–278.

[30] R. Foley and C. Gamble, The ecology of social transitions in human evolution, Philosophical Transactions of the Royal Society B: Biological Sciences, 364 (2009), pp. 3267–3279, https://doi.org/10.1098/rstb.2009.0136.

[31] L. Gabora, Autocatalytic closure in a cognitive system: A tentative scenario for the origin of culture, Psycoloquy, 9 (1998), pp. [adap–org/9901002].

[32] L. Gabora, Conceptual closure: How memories are woven into an interconnected worldview., in Closure: Emergent Organizations and their Dynamics, G. Van de Vijver and J. Chandler, eds., no. 901 in Annual Review Series, Annals of the New York Academy of Sciences, 2000, pp. 42–53, https://doi.org/10.1111/j.1749-6632.2000.tb06264.x.

[33] L. Gabora, Contextual focus: A cognitive explanation for the cultural transition of the Middle/Upper Paleolithic., in Proceedings of the 25th Annual Meeting of the Cognitive Science Society, A. R. and H. D., eds., Lawrence Erlbaum Associates, Hillsdale, NJ, 2003, pp. 432–437.

[34] L. Gabora, Revenge of the ‘neurds’: Characterizing creative thought in terms of the structure and dynamics of human memory., Creativity Research Journal, 22 (2010), pp. 1–13.

[35] L. Gabora, An evolutionary framework for culture: Selectionism versus communal exchange, Physics of Life Reviews, 10 (2013), pp. 117–145, https://doi.org/10.1016/j.plrev.2013.03.006.

[36] L. Gabora, How insight emerges in a distributed, content-addressable memory, in The Cambridge handbook of the neuroscience of creativity, O. Vartanian and J. Jung, eds., MIT Press, Cambridge, MA, 2018, pp. 58–70.

[37] L. Gabora, Reframing convergent and divergent thought for the 21st century, in Proceedings of the 2019 nnual Meeting of the Cognitive Science Society, A. Goel, C. Seifert, and C. Freska, eds., Cognitive Science Society, Austin, TX, 2019, pp. 1794–1800.

[38] L. Gabora and D. Aerts, A model of the emergence and evolution of integrated worldviews, Journal of Mathematical Psychology, 53 (2009), pp. 434–451, https://doi.org/10.1016/j.jmp.2009.06.004.

[39] L. Gabora, S. Leijnen, T. Veloz, and C. Lipo, A non-phylogenetic conceptual network architecture for organizing classes of material artifacts into cultural lineages, in Proceedings of the 33rd annual meeting of the Cognitive Science Society, L. Carlson, C. Hölscher, and T. F. Shipley, eds., Cognitive Science Society, Psychology Press, 2011, pp. 2923–2928.

[40] L. Gabora and C. Smith, Two cognitive transitions underlying the capacity for cultural evolution, Journal of Anthropological Science, 96 (2018), pp. 27–52, https://doi.org/10.4436/jass.96008.

[41] L. Gabora and C. Smith, Exploring the psychological basis for transitions in the archaeological record, in Handbook of cognitive archaeology: Psychology in Prehistory, T. Henley, E. Kardas, and M. Rossano, eds., Routledge / Taylor and Francis, Abingdon, UK, 2019, ch. 12.

[42] L. Gabora and M. Steel, Autocatalytic networks in cognition and the origin of culture, Journal of Theoretical Biology, 431 (2017), pp. 87–95, https://doi.org/10.1016/j.jtbi.2017.07.022.

[43] L. Gabora and M. Steel, Modeling a cognitive transition at the origin of cultural evolution using autocatalytic networks, Cognitive Science, (in press).

[44] X. Gao, W. Huang, X. Ziqiang, M. Zhibang, and J. W. Olsen, 120-150 ka human tooth and ivory engravings from Xinglongdong Cave, Three Gorges Region, South China, Chinese Science Bulletin, 49 (2004), pp. 175–180.

[45] T. L. Griffiths, M. Steyvers, and J. B. Tenenbaum, Topics in semantic representation, Psychological Review, 114 (2007), pp. 211–244, https://doi.org/10.1037/0033-295X.114.2.211.

[46] G. Grimmett and D. Stirzaker, Probability and random processes (3rd ed.), Oxford University Press, 2001.

[47] D. M. Gysi and K. Nowick, Construction, comparison and evolution of networks in life sciences and other disciplines, Journal of the Royal Society Interface, 17 (2020), p. 20190610, https://doi.org/doi.org/10.1098/rsif.2019.0610.

[48] J. Hahn, Kraft und aggression. Die botschaft der eiszeitkunst im Aurignacien Süddeutschlands?, Archaeologica Venatoria, Tübingen, 1986.

[49] J. A. Hampton, Disjunction of natural concepts, Memory and Cognition, 16 (1988), pp. 579–591, https://doi.org/10.3758/BF03197059.

[50] T. Henley, M. J. Rossano, and E. Kardas, Handbook of cognitive archaeology: A psychological framework, Routledge / Taylor and Francis, Abingdon, UK, 2020, https://doi.org/10.4324/9780429488818.

[51] C. Heyes, Enquire within: Cultural evolution and cognitive science, Philosophical Transactions of the Royal Society B: Biological Sciences, 373 (2018), p. 20170051, https://doi.org/10.1098/rstb.2017.0051.

[52] C. J. Holden and R. Mace, Spread of cattle led to the loss of matrilineal descent in africa: a coevolutionary analysis, Proceedings of the Royal Society of London. Series B: Biological Sciences, 270 (2003), pp. 2425–2433, https://doi.org/10.1098/rspb.2003.2535.

[53] W. Hordijk, J. Hein, and M. Steel, Autocatalytic sets and the origin of life, Entropy, 12 (2010), pp. 1733–1742, https://doi.org/10.3390/e12071733.

[54] W. Hordijk, S. A. Kauffman, and M. Steel, Required levels of catalysis for emergence of autocatalytic sets in models of chemical reaction systems, International Journal of Molecular Science, 12 (2011), pp. 3085–3101, https://doi.org/10.3390/ijms12053085.

[55] W. Hordijk and M. Steel, Detecting autocatalytic, self-sustaining sets in chemical reaction systems, Journal of Theoretical Biology, 227 (2004), pp. 451–461, https://doi.org/10.1016/j.jtbi.2003.11.020.

[56] W. Hordijk and M. Steel, Autocatalytic sets and boundaries, J. Syst. Chem., 6:1 (2015).

[57] W. Hordijk and M. Steel, Chasing the tail: The emergence of autocatalytic networks, Biosystems, 152 (2016), pp. 1–10, https://doi.org/10.1016/j.biosystems.2016.12.002.

[58] E. Hovers, S. Lani, O. Bar-Yosef, and B. Vandermeersch, An early case of color symbolism: Ochre use by modern humans in Qafzeh cave, Current Anthropology, 44 (2003), pp. 491–522.

[59] M. W. Howard and M. J. Kahana, A distributed representation of temporal context, Journal of Mathematical Psychology, 46 (2002), pp. 269–299, https://doi.org/10.1006/jmps.2001.1388.

[60] M. N. Jones and D. J. K. Mewhort, Representing word meaning and order information in a composite holographic lexicon, Psychological Review, 114 (2007), pp. 1–37, https://doi.org/10.1037/0033-295X.114.1.1.

[61] P. Kanerva, Hyperdimensional computing: An introduction to computing in distributed representations with high-dimensional random vectors, Cognitive Computation, 1 (2009), pp. 139–159, https://doi.org/10.1007/s12559-009-9009-8.

[62] E. A. Karuza, S. L. Thompson-Schill, and D. S. Bassett, Local patterns to global architectures: influences of network topology on human learning, Trends in Cognitive Sciences, 20 (2016), pp. 629–640.

[63] S. A. Kauffman, Autocatalytic sets of proteins, Journal of Theoretical Biology, 119 (1986), pp. 1–24, https://doi.org/10.3390/ijms12053085.

[64] S. A. Kauffman, The origins of order, Oxford University Press, 1993.

[65] C. Kind, N. Ebinger-Rist, S. Wolf, T. Beutelspacher, and K. Wehrberger, The smile of the lion man. Recent excavations in Stadel cave (Baden-Württemberg, southwestern Germany) and the restoration of the famous upper palaeolithic figurine, Quartär, 61 (2014), pp. 129–145.

[66] M. Kissel and A. Fuentes, ‘behavioral modernity’ as a process, not an event, in the human niche, Time and Mind, 11 (2018), pp. 163–183.

[67] P. J. Kwantes, Using context to build semantics, Psychonomic Bulletin & Review, 12 (2005), pp. 703–710, https://doi.org/10.3758/BF03196761.

[68] P. Lieberman, Language did not spring forth 100,000 years ago, PLoS Biology, 13 (2015), pp. e1002064, doi.org/10.1371/journal.pbio.1002064.

[69] P. M., Wired for culture: The natural history of human cooperation, Penguin, London, UK, 2012.

[70] S. McBrearty and A. Brooks, The revolution that wasn’t: A new interpretation of the origin of modern human behavior, Journal of Human Evolution, 39 (2000), pp. 453–563.

[71] J. M. McClelland, Memory as a constructive process: The parallel distributed processing approach, in The memory process: Neuroscientific and humanistic perspectives, S. Nalbantian, P. M. Matthews, and J. L. McClelland, eds., MIT Press, Cambridge, MA, 2011, pp. 129–155.

[72] J. D. Medaglia, M. E. Lynall, and D. S. Bassett, Cognitive network neuroscience, Journal of Cognitive Neuroscience, 27 (2015), pp. 1471–1491.

[73] S. A. Mednick, The associative basis of the creative process, Psychological Review, 69 (1962), pp. 220–232.

[74] P. Mellars, Going east: New genetic and archaeological perspectives on the modern human colonization of Eurasia, Science, 313 (2006), pp. 796–800.

[75] A. Mesoudi, A. Whiten, and K. N. Laland, Towards a unified science of cultural evolution, Behavioral and Brain Science, 29 (2006), pp. 329–347, https://doi.org/10.1017/S0140525X06009083.

[76] S. Mithen, The prehistory of the mind: A search for the origins of art, religion, and science, Thames and Hudson, London, UK, 1996.

[77] S. Mithen, Creativity in human evolution and prehistory,Routledge, London, UK, 1998.

[78] S. Mithen, Ethnobiology and the evolution of the human mind, Journal of the Royal Anthropological Institute, 12 (2006), pp. S45–S61.

[79] E. Mossel and M. Steel, Random biochemical networks and the probability of self-sustaining autocatalysis, Journal of Theoretical Biology, 233 (2005), pp. 327–336, https://doi.org/10.1016/j.jtbi.2004.10.011.

[80] J. Mulvaney and J. Kamminga, Prehistory of Australia, Smithsonian Institution Scholarly Press, Washington, 1999.

[81] M. Muthukrishna, M. Doebeli, M. Chudek, and J. Henrich, The cultural brain hypothesis: How culture drives brain expansion, sociality, and life history, PLoS Computational Biology, 14 (2018), p. e1006504, https://doi.org/10.1371/journal.pcbi.1006504.

[82] S. Nelson, Diversity of the Upper Palaeolithic Venus figurines and archaeological mythology, Archeological Papers of the American Anthropological Association, 2 (2008), pp. 11–22.

[83] B. A. Nosek, Implicit-explicit relations, Current Directions in Psychological Science, 16 (2007), pp. 65–69.

[84] D. N. Osherson and E. E. Smith, On the adequacy of prototype theory as a theory of concepts, Cognition, 9 (1981), pp. 35–58, https://doi.org/10.1016/0010-0277(81)90013-5.

[85] M. Otte, The management of space during the Paleolithic, Quaternart International, 247 (2012), pp. 212–229.

[86] A. Pike, D. Hoffmann, M. García-Diez, P. Pettitt, J. Alcolea, R. De Balbin, and J. Zilhão, U-series dating of Paleolithic art in 11 caves in Spain, Science, 336 (2012), pp. 1409–1413.

[87] M. Porr, Palaeolithic art as cultural memory: A case study of the Aurignacian art of Southwest Germany, Cambridge Archaeological Journal, 20 (2010), pp. 87–108.

[88] A. Powell, S. Shennan, and M. G. Thomas, Late Pleistocene demography and the appearance of modern human behavior, Science, 324 (2009), pp. 1298–1301.

[89] R. Rappaport, Ritual and religion in the making of humanity, Cambridge University Press, Cambridge, 1999.

[90] E. Rosch, C. B. Mervis, W. D. Gray, D. M. Johnson, and P. Boyes-Braem, Basic objects in natural categories, Cognitive Psychology, 8 (1976), pp. 382–439.

[91] J. M. Smith and E. Szathmary, The major transitions in evolution, Oxford University Press, Oxford, UK, 1997.

[92] P. Sowden, A. Pringle, and L. Gabora, The shifting sands of creative thinking: Connections to dual process theory, Thinking & Reasoning, 21 (2015), pp. 40–60.

[93] M. Steel, W. Hordijk, and J. C. Xavier, Autocatalytic networks in biology: structural theory and algorithms, Journal of the Royal Society Interface, 16 (2019), p. rsif.2018.0808, https://doi.org/10.1098/rsif.2018.0808.

[94] K. Sterelny, *From hominins to humans: how* sapiens *became behaviourally modern*, Philosophical Transactions of the Royal Society of London. Series B, Biological Sciences, 366 (2011), pp. 809–822.

[95] D. Stout, N. Toth, K. Schick, and T. Chaminade, Neural correlates of early stone age toolmaking: technology, language and cognition in human evolution, Philosophical Transactions of the Royal Society B: Biological Sciences, 363 (2008), pp. 1939–1949, https://doi.org/10.1098/rstb.2008.0001.

[96] T. Suddendorf, D. R. Addis, and M. C. Corballis, Mental time travel and the shaping of the human mind, Philosophical Transactions of the Royal Society B: Biological Sciences, 364 (2009), p. 1317, https://doi.org/10.1098/rstb.2008.0301.

[97] I. Tattersall, The origin of the human capacity, American Museum of Natural History, 1998.

[98] J. J. Tehrani and F. Riede, Towards an archaeology of pedagogy: Learning, teaching and the generation of material culture traditions, World Archaeology, 40 (2008), pp. 316–331.

[99] M. Tomasello, A natural history of human thinking, Harvard University Press, Cambridge MA, 2014.

[100] V. Vasas, C. Fernando, M. Santos, S. Kauffman, and E. Szathmáry, Evolution before genes, Biology Direct, 7 (2012).

[101] T. Veloz, L. Gabora, M. Eyjolfson, and D. Aerts, Toward a formal model of the shifting relationship between concepts and contexts during associative thought, in Proceedings of the Fifth International Symposium on Quantum Interaction, D. Song, M. Melucci, I. Frommholz, P. Zhang, L. Wang, and S. Arafat, eds., Springer, Cognitive Science Society, 2011, pp. 25–34, https://doi.org/10.1007/978-3-642-24971-6_4.

[102] T. Veloz, I. Tempkin, and L. Gabora, A conceptual network-based approach to inferring cultural phylogenies, in Proceedings of the 34th annual meeting of the Cognitive Science Society, N. Miyake, D. Peebles, and R. P. Cooper, eds., Cognitive Science Society, Austin TX, 2012, pp. 2487–2492.

[103] A. Whiten, The scope of culture in chimpanzees, humans and ancestral apes, Philosophical Transactions of the Royal Society, Series B., 366 (2011), pp. 997–1007.

[104] A. Whiten, Cultural evolution in animals, Annual Review of Ecology, Evolution, and Systematics, 50 (2019), pp. 1–22.

[105] J. C. Xavier, W. Hordijk, S. Kauffman, S. M., and W. F. Martin, Autocatalytic chemical networks at the origin of metabolism, Proceedings of the Royal Society of London. Series B: Biological Sciences, 287 (2020), p. 20192377.

[106] X. Zhang, D. Wang, and T. Wang, Inspiration or preparation? Explaining creativity in scientific enterprise, in Proceedings of the 25th ACM International on Conference on Information and Knowledge Management, ACM, 2016, pp. 741–750.

